# Enhancing vs. inhibiting semantic performance with repetitive transcranial magnetic stimulation over the anterior temporal lobe: frequency- and task- specific effects

**DOI:** 10.1101/2019.12.16.878256

**Authors:** JeYoung Jung, Matthew A. Lambon Ralph

## Abstract

Accumulating, converging evidence indicates that the anterior temporal lobe (ATL) appears to be the transmodal hub for semantic representation. A series of repetitive transcranial magnetic stimulation (rTMS) investigations utilizing the ‘virtual lesion’ approach have established the brain-behavioural relationship between the ATL and semantic processing by demonstrating that inhibitory rTMS over the ATL induced impairments in semantic performance in healthy individuals. However, a growing body of rTMS studies suggest that rTMS might also be a tool for cognitive enhancement and rehabilitation, though there has been no previous exploration in semantic cognition. Here, we explored a potential role of rTMS in enhancing and inhibiting semantic performance with contrastive rTMS protocols (1Hz vs. 20Hz) by controlling practice effects. Our results demonstrated that it is possible to modulate semantic performance positively or negatively depending on the ATL stimulation frequency: 20Hz rTMS was optimal for facilitating cortical processing (faster RT in a semantic task) contrasting with diminished semantic performance after 1Hz rTMS. In addition to cementing the importance of the ATL to semantic representation, our findings suggest that 20Hz rTMS leads to semantic enhancement in healthy individuals and potentially could be used for patients with semantic impairments as a therapeutic tool.

## Introduction

Concepts and meaning are fundamental components of human cognition. We use this knowledge every day to recognise objects in our environment, to anticipate how they will behave and interact with each other and, use them to perform functions, to generate expectations for situations, and to interpret language. Converging evidence from neuropsychological and neuroscientific studies indicates that the anterior temporal lobe (ATL) is a central area serving as a representational hub interacting with distributed modality-specific ‘spoke’ regions in order to form coherent and generalizable concepts (for a review, see Patterson K et al. 2007; Lambon Ralph MA et al. 2010; Lambon Ralph MA 2014; Ralph MA et al. 2017). Semantic dementia (SD; the temporal lobe variant of frontotemporal dementia) is a most striking example supporting this hypothesis – SD patients with atrophy centred in the ATL exhibit a selective semantic impairment in both verbal and non-verbal domains (Bozeat S et al. 2000; Mummery CJ et al. 2000; Hodges JR and K Patterson 2007). This hypothesis was subsequently supported and extended by recent investigations using intracranial recording (Abel TJ et al. 2015; Shimotake A et al. 2015; Chen Y et al. 2016), magnetoencephalography (MEG) (Clarke A et al. 2011; Mollo G et al. 2017), and functional magnetic resonance imaging (fMRI) (Visser M et al. 2010; Peelen MV and A Caramazza 2012; Coutanche MN and SL Thompson-Schill 2015; Murphy C et al. 2017).

A crucial form of convergent evidence for the causal role of the ATL in semantic representation came through a series of experiments with healthy participants using repetitive transcranial magnetic stimulation (rTMS) demonstrating that rTMS over the ATL causes transient impairments in various semantic tasks (Pobric G et al. 2007; Lambon Ralph MA et al. 2009; Pobric G et al. 2009; Binney RJ et al. 2010; Pobric G, E Jefferies and MA Lambon Ralph 2010; Pobric G, E Jefferies and MA Ralph 2010; Jackson RL et al. 2015; Jung J and MA Lambon Ralph 2016). Although this ‘virtual lesion’ rTMS approach has been useful to verify brain-behaviour relationships, several studies have shown enhancement in cognitive performance, suggesting that rTMS is also capable of facilitating cortical activity at the site of simulation (for a review, see Vallar G and N Bolognini 2011), depending on the type and frequency of stimulation (Miniussi C and PM Rossini 2011). Furthermore, there is a growing interest in the possibility of utilising TMS in cognitive rehabilitation (Rossi S and PM Rossini 2004; Miniussi C and PM Rossini 2011) and enhancement (Luber B and SH Lisanby 2014). In this study, therefore, we tested various rTMS protocols for enhancing vs diminishing semantic processing.

Converging evidence indicates that rTMS below 1Hz reduces cortical excitability at the target region (Chen R et al. 1997), whereas high frequency (HF) rTMS (typically between 5 and 20Hz) can increase it (Pascual-Leone A et al. 1994). With this ability, rTMS has been widely used to manipulate cortical processing and to examine the resultant changes in cognitive performance. Studies reported TMS-induced performance enhancements have employed various TMS protocols including single pulse, theta burst, paired pulse, and rTMS at both low and high frequencies and cognitive tasks (for a review, see Miniussi C and PM Rossini 2011; Luber B and SH Lisanby 2014). Cortical processing are affected differently by these various forms of TMS: some disrupting processing through the addition of neural noise; briefly inhibiting or facilitating activity; whilst others modulate cortical excitability up or down for periods beyond the stimulation. Luber and Lisanby (2014) suggested that there are three mechanisms underlying TMS enhancement effects: nonspecific effects of TMS, disruption of competing processing (i.e., addition-by-subtraction), and direct modulation of TMS to task-related cortex. Non-specific TMS effects are ‘side effects’ of the stimulation (e.g., increased alertness following the ‘click’ sound or tactile sensation). These peripheral sensations can cause intersensory facilitation that contributes to performance enhancements (Terao Y et al. 1997). An example of ‘addition-by-subtraction’ is that inhibitory 10mins 1Hz rTMS applied to the right posterior parietal cortex (involved in directing attention to salient stimuli) improved reaction time (RT) in a visual search task when there are attention-capturing distracters (Hodsoll J et al. 2009). Many HF rTMS studies (between 5 and 20Hz and iTBS; intermittent theta burst stimulation) have successfully produced performance enhancements via direct TMS modulation to task-related cortex (Ragert P et al. 2003; Wagner M et al. 2006; Boyd LA and MA Linsdell 2009; Hwang JH et al. 2010; Ahn HM et al. 2013; Hoy KE et al. 2016). The associated, prolonged facilitatory effects are thought to be based on long term potentiation (LTP) (Bliss TV and T Lomo 1973). Here, we attempted to achieve semantic performance enhancements via direct modulation by stimulating the ATL with HF rTMS, and contrasted this to the transient inhibition induced by low-frequency stimulation to the same region, in the same participants.

We also tackled a crucial, additional cognitive factor that is rarely addressed in rTMS enhancements studies, namely practice effects (performance enhanced through repeated exposure to the test procedure and stimuli). Many rTMS enhancements studies have sought to measure changes in behavioural performance by comparing two sessions, before and after stimulation. However, the effects of practice at such brief test-retest intervals have not been considered as a significant factor in TMS literature. Practice effects on cognitive performance vary according to the difficulty of the task (stronger for more difficult tasks), the length of the test-retest interval (practice effects diminish over time), the individual’s ability at the time of testing (stronger practice effects for weaker participants) and can be reduced through the use of alternative forms or stimuli sets (Benedict RH and DJ Zgaljardic 1998; Basso MR et al. 1999; Dikmen SS et al. 1999) though practice-related performance improvement can still persist (Kay G 1991). Studies investigating practice effects on cognitive test performance suggest that at least two assessments need to be conducted before performance stabilizes and there are no practice effects on simple tasks with longer (i.e., one week) test-retest intervals (Collie A et al. 2003; Falleti MG et al. 2006). Accordingly in the current study, we asked participants to perform a thorough task familiarization procedure in order to establish a ‘stable baseline’ prior to stimulation and thus minimize practice-related improvement in the experiment itself (see Materials and Methods).

Given that there is no consensus on a set rTMS protocol for inducing facilitatory effects, we first evaluated various rTMS protocols that have produced performance enhancements on higher cognition functions in healthy participants. The four selected protocols (5, 10, and 20Hz, and iTBS) have showed facilitatory effects on working memory, learning and executive functions in previous investigations (see the Materials and Methods). In the pilot study, we applied rTMS at four HF over the left ventrolateral ATL (vATL) in healthy participants to identify which protocols induce semantic performance enhancements. Early rTMS studies stimulated at the lateral ATL, 10mm posterior to the tip of the temporal pole on the middle temporal gyrus on the basis that this fell into the area of atrophy observed in SD patients (Pobric G *et al*. 2007; Lambon Ralph MA *et al*. 2009; Pobric G, E Jefferies and MA Lambon Ralph 2010). Recent convergent evidence from patient, fMRI and cortical electrode has shown that the vATL region appears to be the centre point of hub with strong multimodal and omni-category responses (Rice GE et al. 2015; Lambon Ralph MA et al. 2017). Also, our recent rTMS-fMRI combined study demonstrated that stimulating the vATL decreased regional activity at the target site and the ventromedial ATL, and induced slowed semantic performance (Jung J and MA Lambon Ralph 2016). Therefore, we chose the vATL as the target site in this study. Having selected 20Hz rTMS from the pilot study, we investigated its effects on semantic performance in comparison to an opposing, inhibitory stimulation (1 Hz) and sham stimulation.

Before and after the stimulation, participants performed an object category judgement task and a pattern matching task serving as a control task. We hypothesized that 20Hz rTMS over the vATL would produce semantic performance enhancements (faster RT) in comparison to no-stimulation (before the stimulation) and sham stimulation, whilst 1Hz rTMS would show the opposite effects on semantic performance. In addition, we expected that semantic processing should be preferentially modulated by both 20Hz and 1Hz rTMS over the left vATL and not the control site.

## Materials and Methods

### Participants

Twenty five healthy participants participated in this study; four participants (3 females, mean age, 23.1 ± 4.2 years) participated in the pilot study in order to determine an optimal rTMS protocol to induce facilitatory effects on semantic processing, and twenty one in the main experiment (7 females, mean age, 22 ± 3.1 years). The sample size was calculated based on a previous study (Jung J and MA Lambon Ralph 2016), which used a 2 (TMS to ATL vs. control site) ×2 (semantic vs. control task) within subject design. These previous data (collected in N=23 participants) indicated that to achieve α=0.05, power=80% for the critical interaction between TMS site and task then N≥18 were required. All participants were native English speakers with right-handed assessed by the Edinburgh Inventory for Handedness (Oldfield RC 1971). They received a detailed explanation of the study and gave written informed consent prior to the experiment. The experiment was approved by the local ethics committee.

### Experimental Design and Procedure

In the pilot study, four participants performed an object category judgment task and a number judgement task as a control task. The stimuli for the category judgment task were from the *Levels of Familiarity, Typicality, and Specificity (LOFTS)* semantic battery (Rogers TT et al. 2015). The 120 items probe semantic knowledge at the subordinate level and cover a variety of categories, including animals, vehicles, tools, foods, and plants (Table S1). Participants were asked to indicate which of two categories was appropriate for a target object (e.g., target: collie, choice 1: dog, choice 2: car). In each trial, three words were presented on the screen, a target on the top and 2 choices at the bottom (Fig. 1A). In the number judgement, control task participants judged whether a two-digit number was even or odd. Participants performed the tasks before and after the stimulation. Each task had 120 trials, a trial started with 500ms fixation and the stimuli were presented until the participant’s response or a maximum of 5000ms. E-prime software (Psychology Software Tools Inc., Pittsburgh, PA, USA) was used to display stimuli and to record responses.

**Figure 1.**
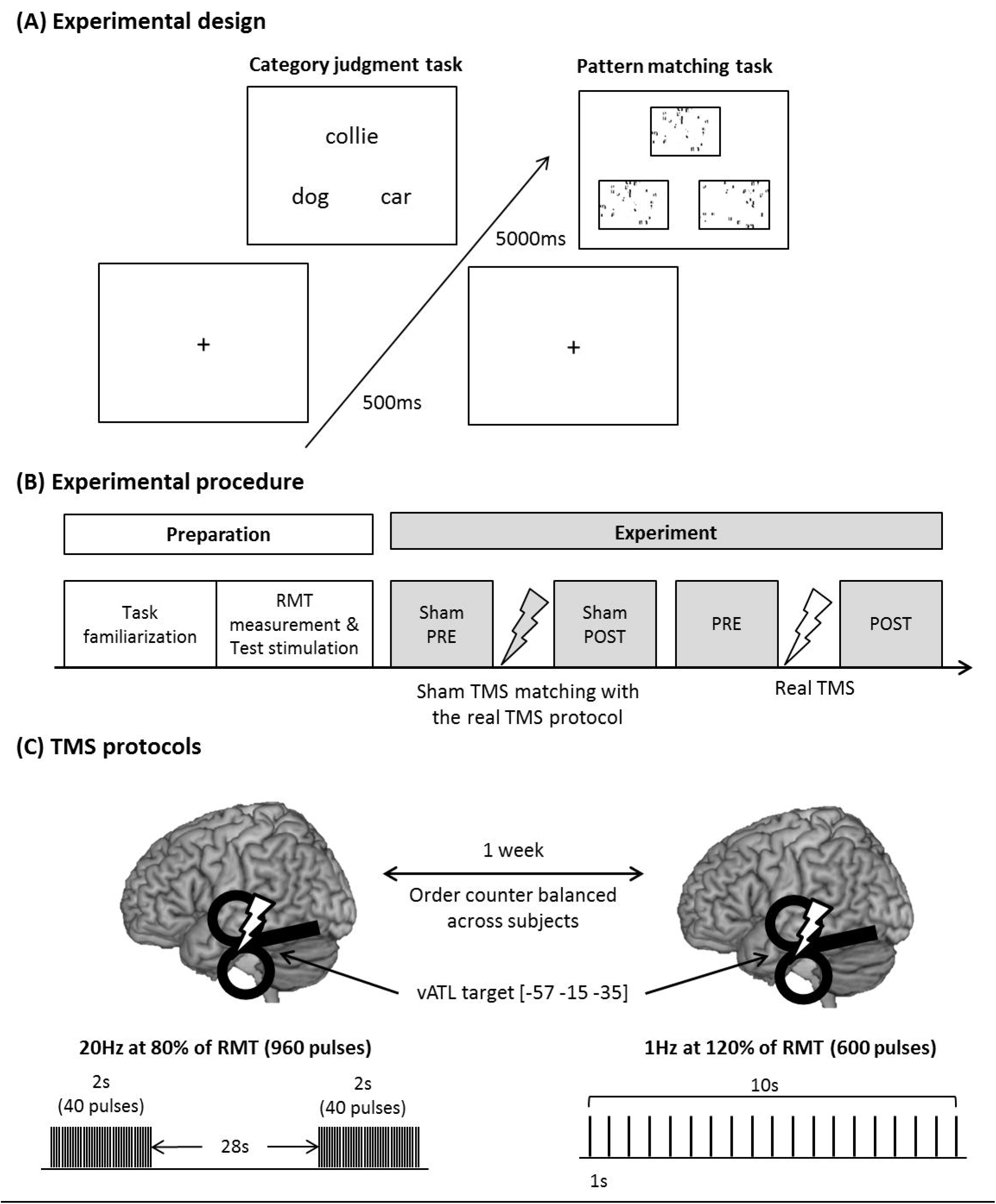
(A) Experimental design. Each trial starts with a fixation followed by stimuli, which have 3 items, a target on the top and 2 choices at the bottom. (B) Experimental procedure. Experimental preparation includes the task familiarization and RMT measurement/test stimulation for following the experiment. During the task familiarization, participants performed 120 trials of each task in order to saturate their task performance thereby minimizing practice effects. During the test stimulation, 100 pulses of TMS matched with the following TMS protocol were delivered over the occipital pole so as to prevent non-specific TMS effects. In the experiment, 4 task sessions were conducted before and after sham and real TMS stimulation. (C) TMS protocols. 2 TMS protocols were employed: 20Hz for the facilitatory effects and 1Hz for the inhibitory effects. Each protocol was delivered on different days with a week gap at least. The stimulation was applied at the ventrolateral anterior temporal lobe.

In the main experiment, twenty-one participants performed the same category judgement task from the pilot study and a pattern matching task as a control task. As the number judgement task used in the pilot study proved not to be well matched with the category task in terms of difficulty (about 110ms faster than the category task, see Fig. S1B), we employed a pattern matching task from previous TMS and fMRI studies (Pobric G *et al*. 2007; Visser M et al. 2012) which provided a better match to the categorisation task in terms of general difficulty. In this pattern matching task, participants were asked to select which of two patterns was identical to a target pattern (Fig. 1A). In order to minimize the practice effects found in the pilot study (Fig. S2), participants performed a task familiarization procedure in which they completed 120 trials of each task prior to the main experiment. Participants then performed the main tasks before and after sham and real TMS (Fig. 1B). The task included 60 trials in the main experiments. E-prime software (Psychology Software Tools Inc., Pittsburgh, PA, USA) was used to display stimuli and to record responses. The experimental design and procedure is summarized in Fig. 1.

### Anatomical MRI Acquisition

A high-resolution T1-weighted anatomical image collected on a 3T Philips MR Achieva scanner was obtained for all participants to guide a target site. The structural image was acquired using a 3D MPRAGE pulse sequence with 200 slices, in planed resolution 0.94 × 0.94, slice thinkness 0.9mm, TR = 8.4ms, and TE = 3.9ms.

### Transcranial Magnetic Stimulation (TMS)

A MagStim Super Rapid stimulator (The MagStim Company, Whitland, UK) was used to deliver stimulation with a figure of eight coil (70mm). Resting motor threshold (RMT) was defined as a minimal intensity of stimulation inducing motor evoked potentials in the contralateral first dorsal interosseous muscle of the left motor cortex in at least 5 of 10 stimulations. The average RMT intensity was 60.6% ± 7.4 in the pilot and 60.7% ± 7.2 in the experiment.

The target site [MNI: Montreal Neurological Institute, −57 −15 −35] was selected from previous fMRI and TMS studies (Visser M *et al*. 2012; Jung J and MA Lambon Ralph 2016). The coordinate was located on the ventrolateral ATL (Fig. 1C) and transformed to each participant’s native space. Statistical Parametric Mapping software (SPM8, Wellcome Trust Centre for Neuroimaging, London, UK) was used to normalize participants’ MRI scan against the MNI template and to convert the target coordinate to the untransformed individual native space coordinate using the inverse of each resulting transformation. These native space coordinates guided the frameless stereotaxy, via a Brainsight TMS-MRI co-registration system (Rogue Research, Montreal, Canada).

### TMS protocol

In order to determine an optimal rTMS protocol, we tested four protocols that have shown facilitatory effects in previous studies: 5Hz, 10Hz, 20Hz, and iTBS. We selected these protocols on the basis that a single session application had induced facilitatory effects on cognitive behaviours or motor evoked potentials. 5Hz protocol had 2 blocks of 9 trains of 10s stimulation repeated every 20s (total 900 pulses) (Sole-Padulles C et al. 2006). 10Hz protocol had 3 blocks of 15 trains of 2s stimulation repeated every 12s (total 900 pulses) (Ahn HM *et al*. 2013). 20Hz stimulation consisted of 3 blocks of 8 trains of 2s stimulation repeated every 28s (total 960 pulses) (Wagner M *et al*. 2006). iTBS (intermittent theta-burst stimulation) had 3 pulses of stimulation given at 50Hz, a 2s train of TBS repeated every 10s for 190s (total 600 pulses) (Huang YZ et al. 2005; Hoy KE *et al*. 2016). As a control, we used sham stimulation in which one wing of a figure-eight coil was in contact with the target site, but at a 90° tilt from tangential (Lisanby SH et al. 2001). All protocols were delivered with 80% of RMT for each individual.

In the pilot study, participants took part in 5 TMS sessions on different days. The order of protocols was counterbalanced across the participants. We conducted paired t-test between four facilitatory protocols and sham. The results demonstrated that 20Hz stimulation induced marginally significant facilitatory effects on semantic processing (Wilcoxon signed-rank test: Z = −1.5, p = 0.07) (Fig. S1C). Therefore, we selected 20Hz protocol for the main experiment.

In the main experiment, we contrasted 20Hz and 1Hz rTMS, and compared these to each other as well as sham to stimulation. 1Hz rTMS to the ATL (total 600 pulses, 120% RMT) has demonstrated inhibitory effects in semantic processing (Pobric G *et al*. 2007; Pobric G *et al*. 2009; Pobric G, E Jefferies and MA Lambon Ralph 2010; Pobric G, E Jefferies and MA Ralph 2010). Here, we expected to replicate the same 1Hz rTMS effect whilst finding the opposite effect for 20Hz stimulation. Sham TMS was delivered with the same protocol of the real TMS on the day of experiment. During the stimulation, participants were asked to be relaxed with closed eyes. Each session was conducted at the same time on different days with a week gap between sessions (Fig. 1C). In order to control for non-specific TMS effects, we delivered 60 pulses of stimulation (test stimulation, matched with the subsequent real TMS protocol) over the occipital pole prior to the experiment, which was localised using the international 10-20 system. Occipital pole is a common control site for TMS studies and previous studies have demonstrate that this site successfully served as the control site and did not influence behavioural performance and neural changes in either semantic or control (visual) tasks (Pobric G *et al*. 2007; Lambon Ralph MA *et al*. 2009; Pobric G, E Jefferies and MA Ralph 2010; Jung J and MA Lambon Ralph 2016). After the experiment, we asked participants whether they could distinguish between sham and real TMS. All participants reported that the sham stimulation was a real stimulation, felt weaker than the real TMS stimulation. There were no complaints about discomfort associated with HF rTMS over the vATL.

## Results

The participants’ performance on the semantic task (category judgment) and the control task (pattern matching) was compared following 20Hz and 1Hz vATL rTMS. Reaction time (RT) was examined using a repeated measures analysis of variance (ANOVA) with protocol (20Hz vs. 1Hz), task (category judgement vs. pattern matching), and TMS (Pre vs. Post) as within-subject factors. There were a significant main effect of TMS (F_1, 20_ = 5.66, p < 0.05) and interactions between the protocol and TMS (F_1, 20_ = 6.29, p < 0.05) and between the protocol, task, and TMS (F_1, 20_ = 7.24, p < 0.05). The other main effects and interactions did not reach the significance level (Fs < 0.57, ps > 0.46). Post hoc paired t-tests evaluating the interactions revealed that RT for the category judgements was slower after the 1Hz stimulation (t = −2.09, p = 0.05) and faster after the 20Hz stimulation (t = 4.37, p < 0.001) (Fig. 2A Left). Accuracy rates were high (PRE-session: category task 92%, pattern matching 95%).

**Figure 2.**
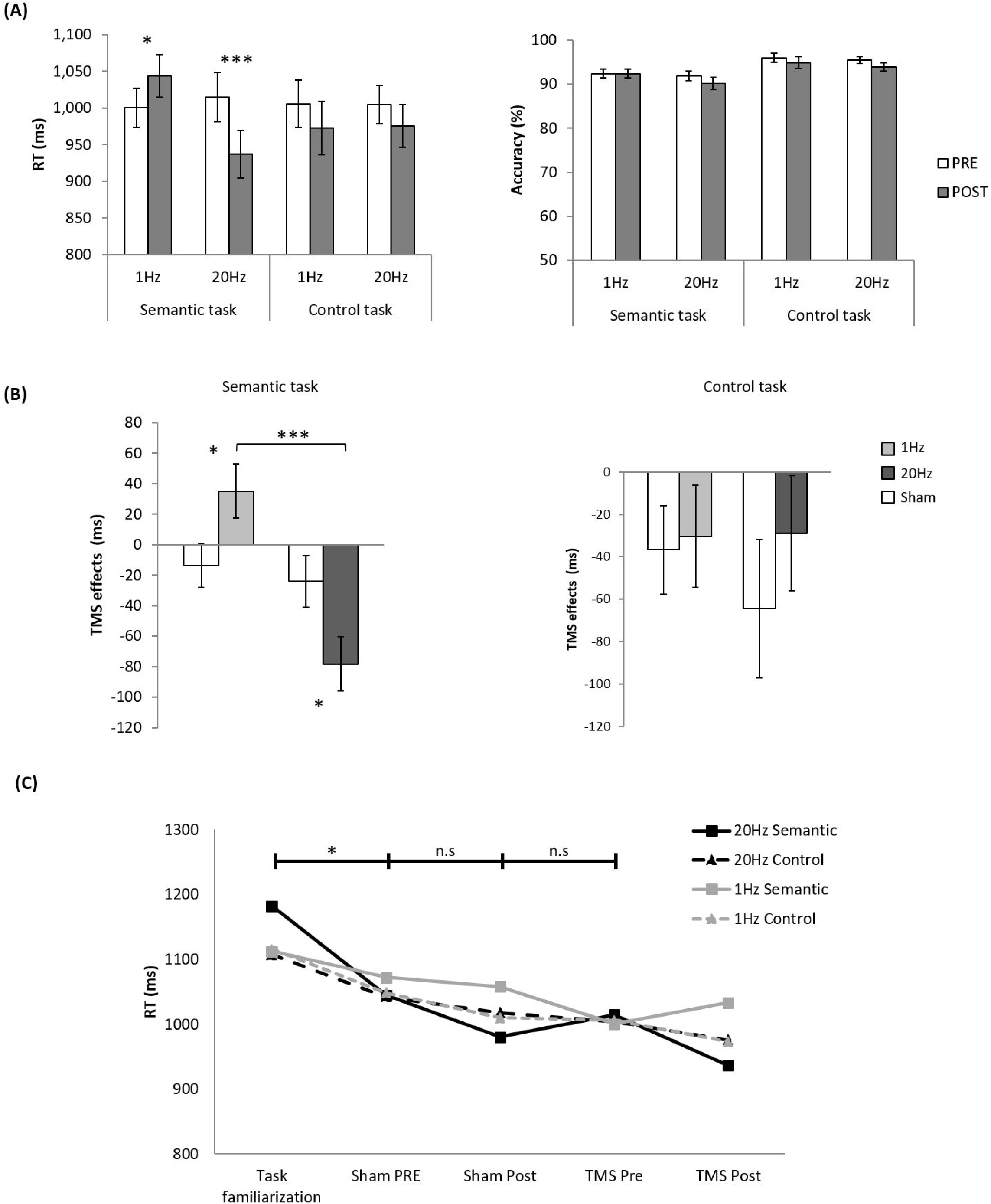
The results of the experiment. (A) The averaged reaction time (RT) and accuracy (%) for the semantic (category judgement) and control (pattern matching) tasks before (PRE) and after (POST) stimulation. White bars represent the PRE sessions and grey bars, the POST sessions. (B) TMS effects in the semantic and control task. TMS effects were computed by subtracting RT in PRE from POST TMS sessions. White bars represent the matched sham stimulation. Light grey bar indicates 1Hz stimulation and dark grey bar, 20Hz stimulation. Error bars represent standard errors. (C) Practice effects. Squares with solid lines indicate semantic task performance, whereas triangles with dotted lines, control task performance. Black colour represents 20Hz stimulation and grey represents 1Hz, control stimulation. * p < 0.05, *** p ≤ 0.001

Participants made less errors in the pattern matching than category judgment task (F_1, 20_ = 25.61, p < 0.001) and showed a tendency of increasing errors after the stimulation (F_1, 20_ = 4.13, p = 0.06) (Fig. 2A Right). However, there was no effect of protocol or an interaction (Fs < 1.8, ps > 0.20). Figure 2 summarizes the results.

As noted in the Introduction, TMS-induced facilitatory effects need to be verified over and above practice effects. Accordingly, we expected that the facilitatory effect induced by TMS should be bigger than practice effects. To test this hypothesis, we calculated the TMS effects (RT difference between pre and post session: POSTPRE) and compared them with the RT changes caused by sham stimulation. The results are summarized in Figure 2B. Planned paired t-tests were conducted between rTMS protocols and the matched sham stimulation. We found that the category judgment times were significantly slower after 1Hz stimulation (t = −2.29, p < 0.05) and faster after 20Hz stimulation (t = 2.14, p < 0.05) compared to the sham stimulation. There was a significant difference between 20Hz and 1Hz stimulation (t = 3.91, p = 0.001). As expected, there was no RT differences for the pattern matching task (ps> 0.22) (Fig. 2B).

Sham stimulation performance was examined using the same analysis. In RT, there was a significant main effect of TMS (F_1, 20_ = 13.69, p < 0.001), but no effect of other factors and interactions (Fs < 3.16, ps > 0.09). The effect of TMS reflects practice effects, i.e., a decrease in RT after any stimulation protocols regardless of tasks. In accuracy, there was a significant main effect of task (F_1, 20_ = 46.49, p < 0.001), but no effect of other factors and interactions (Fs < 3.15, ps > 0.10). Again, the accuracy rates were high in both tasks (Pre-session: category judgement 91%, pattern matching 95%). The results are summarized in Supplementary Figure S3.

To evaluate practice effects between sessions (task familiarization, sham PRE and POST, and TMS PRE), we conducted a one-way ANOVA for each task. Figure 2C summarises the results. In RT, we found practice effects (20Hz semantic task: F_3, 76_ = 2.72, p = 0.05, 20Hz control task: F_3, 76_ = 1.98, p = 0.10, 1Hz semantic task: F_3, 76_ = 2.94, p < 0.05, 1Hz control task: F_3, 76_ = 5.12, p < 0.01). Post hoc paired t-tests demonstrated that there was a significant difference between the first and second sessions (task familiarization and sham PRE) and no difference between the second and third sessions and between the third and fourth sessions. There was no significant practice effect in accuracy (Fs < 0.88, ps > 0.22).

## Discussion

There is a growing interest in TMS as a tool for cognitive enhancement and rehabilitation, in addition to the more established use for evaluating the effect of TMS-induced cortical inhibition on cognitive function. Many studies employing various TMS paradigms have showed a facilitatory effect on motor and cognitive functions such as memory, language, executive functions (Rossi S and PM Rossini 2004; Luber B and SH Lisanby 2014). To date, however, TMS-induced enhancements have not been explored in the semantic domain. Depending on the TMS frequency, we found that semantic performance could be improved (20Hz) or impeded (1Hz) in the same participants. We also confirmed that the enhancing effect of HF vATL rTMS was not due to uncontrolled nuisance effects, including practice effects. Our results suggest that 20Hz vATL rTMS is an optimal protocol for facilitating cortical processing during semantic categorisation and thus might be a beneficial intervention for semantic enhancement in healthy individuals and rehabilitation in patients.

To our best knowledge, this is the first study to demonstrate (a) opposing effects of vATL TMS on semantic performance in the same participants dependent on the frequency used; and (b) semantic enhancements in healthy participants by stimulating the vATL with a facilitatory protocol. The ability to both inhibit and improve semantic performance provides further strong evidence for the role of the ATL as a transmodal hub for semantic representation. The demonstration of enhanced semantic performance is a crucial addition to the multiply replicated effect of inhibitory rTMS over the ATL (1Hz rTMS and cTBS; continuous theta burst stimulation) during various semantic tasks (Pobric G *et al*. 2007; Lambon Ralph MA *et al*. 2009; Pobric G *et al*. 2009; Pobric G, E Jefferies and MA Lambon Ralph 2010; Pobric G, E Jefferies and MA Ralph 2010; Jung J and MA Lambon Ralph 2016). These past studies showed that inhibitory ATL rTMS induced slowed RT during semantic processing. We note here that a recent study employing a shorten version of cTBS (20s, 300 pulses) over the temporal pole showed a partial facilitatory effect on semantic processing (Bonni S et al. 2015). cTBS over the right temporal pole was only found to improve RT during picture-based semantic processing in comparison to control stimulation (vertex) whilst there was no TMS effects after the left temporal pole stimulation in both picture and word semantic association tasks. These findings are inconsistent with the many previous ATL rTMS studies but it should be noted that (a) they delivered only half of the typical dose for the cTBS protocol (40s, 600 pulses) (Huang YZ *et al*. 2005), and (b) that inhibitory rTMS protocols may produce behavioural facilitation on higher cognitive domains due to the alerting effect of TMS (Vallar G and N Bolognini 2011) or uncontrolled practice effects. In comparison, our study was able to establish and evaluate optimal protocols for inhibitory and excitatory TMS for vATL-related semantic processing, and to demonstrate these effects over and above any alerting or practice.

The underlying mechanisms of rTMS effects are not fully understood but studies have showed that the most likely mechanisms relate to changes in synaptic transmission between neurons including long-term potentiation (LTP) and long-term depression (LTD) (for a review, see Pell GS et al. 2011). Low frequency rTMS (~1Hz and cTBS) can induce a suppression of cortical excitability whereas HF rTMS (5~20Hz and iTBS) potentiate it (Fitzgerald PB et al. 2006). These effects have been found to depend on both γ-aminobutyric acid (GABA) and glutamate system (NMDA and AMPA receptors) activity (Funke K and A Benali 2011; Lenz M et al. 2016). Recent animal studies have unveiled the underlying cellular and molecular mechanisms of rTMS *in vivo* and *in vitro* (i.e., LTP/LTD, spike-timing-dependent plasticity in excitatory and inhibitory synapses, intrinsic cellular plasticity, metaplastcity, and structural plasticity) (Muller-Dahlhaus F and A Vlachos 2013). HF rTMS leads to long-lasting structural and functional changes in excitatory and inhibitory postsynapses. For example, 10Hz rTMS (9 trains of 100 pulses with an interval 30s) applied over the slice cultures of rat CA1 pyramidal neurons, increased excitatory post synaptic transmission, dendritic spin size in excitatory synapses (Vlachos A et al. 2012) and decreased inhibitory synaptic transmission accompanied with the reduction in related inhibitory receptor properties (GABA) (Lenz M *et al*. 2016). The observed facilitatory TMS effects in the current study may be attributed to these molecular mechanisms underlying synaptic plasticity.

Accumulating evidence indicates that rTMS causes not only local changes, but also modulation of remote but functionally connected brain regions (O’Shea J et al. 2007; Bestmann S et al. 2008). Thus, the long-term after effects can be attributed to activity changes in a given network rather than a local excitation or inhibition of an individual region alone. Our previous studies employing rTMS combined with fMRI showed that cTBS over the left vATL reduced activity in the target site as well as upregulation in the contralateral vATL and increased inter-ATL functional connectivity, which contributed to semantic performance after the stimulation (Binney RJ and MA Lambon Ralph 2014; Jung J and MA Lambon Ralph 2016). Future studies that combine excitatory 20Hz vATL rTMS with fMRI will be able to explore whether its positive effects reflect changes in the stimulated region and/or networklevel modulations.

It is generally agreed that the effect of rTMS is primarily determined by the specific combination of stimulation frequency and intensity (Wassermann EM et al. 2008). However, many other factors can influence rTMS effects such as history of synaptic activity, attention, time of day, and age (Ridding MC and U Ziemann 2010). In this study, we tried to control these nuisance factors in order to delineate task-specific facilitatory TMS effects on higher cognition. The pilot study illustrated that there were huge practice effects (the baseline of 5 sessions showed gradual reduction in RT according to the order of the session, see Fig. S2) and, even in the sham stimulation, practice effects were prominent (Fig. S1). Thus, we included a substantial task familiarization procedure to minimize practice effects and to stabilize task performance prior to the main experiment, and each rTMS protocol was conducted at least a week apart. As a result, non-specific improvements in task performance were saturated prior to the TMS sessions (Fig. 2A) allowing us to be much more more sensitive to TMS-induced changes in semantic performance.

In conclusion, we identified an optimal vATL rTMS protocol, which can induce either semantic enhancement or inhibition in healthy individuals. Our results demonstrated that 20Hz rTMS over the left vATL induced semantic performance enhancement – faster RT during a category judgment task, contrasting with 1Hz rTMS – slower semantic decisions. Our findings not only add important new causal evidence for the vATL as the major hub for semantic representation but also indicate that HF rTMS could be a potential therapeutic tool in cognitive rehabilitation for patients with semantic impairments.

## Acknowledgments

This research was supported by a Beacon Anne McLaren Research Fellowship (University of Nottingham) to JJ and an Advanced ERC award (GAP: 670428 - BRAIN2MIND_NEUROCOMP) and MRC programme grant (MR/R023883/1) to MALR.

## References

Abel TJ, Rhone AE, Nourski KV, Kawasaki H, Oya H, Griffiths TD, Howard MA, 3rd, Tranel D. 2015. Direct physiologic evidence of a heteromodal convergence region for proper naming in human left anterior temporal lobe. J Neurosci. 35:1513–1520.

Ahn HM, Kim SE, Kim SH. 2013. The effects of high-frequency rTMS over the left dorsolateral prefrontal cortex on reward responsiveness. Brain Stimul. 6:310–314.

Basso MR, Bornstein RA, Lang JM. 1999. Practice effects on commonly used measures of executive function across twelve months. Clin Neuropsychol. 13:283–292.

Benedict RH, Zgaljardic DJ. 1998. Practice effects during repeated administrations of memory tests with and without alternate forms. J Clin Exp Neuropsychol. 20:339–352.

Bestmann S, Ruff CC, Blankenburg F, Weiskopf N, Driver J, Rothwell JC. 2008. Mapping causal interregional influences with concurrent TMS-fMRI. Exp Brain Res. 191:383–402.

Binney RJ, Embleton KV, Jefferies E, Parker GJ, Ralph MA. 2010. The ventral and inferolateral aspects of the anterior temporal lobe are crucial in semantic memory: evidence from a novel direct comparison of distortion-corrected fMRI, rTMS, and semantic dementia. Cereb Cortex. 20:2728–2738.

Binney RJ, Lambon Ralph MA. 2014. Using a combination of fMRI and anterior temporal lobe rTMS to measure intrinsic and induced activation changes across the semantic cognition network. Neuropsychologia.

Bliss TV, Lomo T. 1973. Long-lasting potentiation of synaptic transmission in the dentate area of the anaesthetized rabbit following stimulation of the perforant path. J Physiol. 232:331–356.

Bonni S, Koch G, Miniussi C, Bassi MS, Caltagirone C, Gainotti G. 2015. Role of the anterior temporal lobes in semantic representations: Paradoxical results of a cTBS study. Neuropsychologia. 76:163–169.

Boyd LA, Linsdell MA. 2009. Excitatory repetitive transcranial magnetic stimulation to left dorsal premotor cortex enhances motor consolidation of new skills. Bmc Neurosci. 10.

Bozeat S, Lambon Ralph MA, Patterson K, Garrard P, Hodges JR. 2000. Non-verbal semantic impairment in semantic dementia. Neuropsychologia. 38:1207–1215.

Chen R, Classen J, Gerloff C, Celnik P, Wassermann EM, Hallett M, Cohen LG. 1997. Depression of motor cortex excitability by low-frequency transcranial magnetic stimulation. Neurology. 48:1398–1403.

Chen Y, Shimotake A, Matsumoto R, Kunieda T, Kikuchi T, Miyamoto S, Fukuyama H, Takahashi R, Ikeda A, Lambon Ralph MA. 2016. The ‘when’ and ‘where’ of semantic coding in the anterior temporal lobe: Temporal representational similarity analysis of electrocorticogram data. Cortex. 79:1–13.

Clarke A, Taylor KI, Tyler LK. 2011. The evolution of meaning: spatio-temporal dynamics of visual object recognition. J Cogn Neurosci. 23:1887–1899.

Collie A, Maruff P, Darby DG, McStephen M. 2003. The effects of practice on the cognitive test performance of neurologically normal individuals assessed at brief test-retest intervals. J Int Neuropsych Soc. 9:419–428.

Coutanche MN, Thompson-Schill SL. 2015. Creating Concepts from Converging Features in Human Cortex. Cereb Cortex. 25:2584–2593.

Dikmen SS, Heaton RK, Grant I, Temkin NR. 1999. Test-retest reliability and practice effects of Expanded Halstead-Reitan neuropsychological test battery. J Int Neuropsych Soc. 5:346–356.

Falleti MG, Maruff P, Collie A, Darby DG. 2006. Practice effects associated with the repeated assessment of cognitive function using the CogState battery at 10-minute, one week and one month test-retest intervals. J Clin Exp Neuropsyc. 28:1095–1112.

Fitzgerald PB, Fountain S, Daskalakis ZJ. 2006. A comprehensive review of the effects of rTMS on motor cortical excitability and inhibition. Clin Neurophysiol. 117:2584–2596.

Funke K, Benali A. 2011. Modulation of cortical inhibition by rTMS - findings obtained from animal models. J Physiol. 589:4423–4435.

Hodges JR, Patterson K. 2007. Semantic dementia: a unique clinicopathological syndrome. Lancet Neurol. 6:1004–1014.

Hodsoll J, Mevorach C, Humphreys GW. 2009. Driven to less distraction: rTMS of the right parietal cortex reduces attentional capture in visual search. Cereb Cortex. 19:106–114.

Hoy KE, Bailey N, Michael M, Fitzgibbon B, Rogasch NC, Saeki T, Fitzgerald PB. 2016. Enhancement of Working Memory and Task-Related Oscillatory Activity Following Intermittent Theta Burst Stimulation in Healthy Controls. Cereb Cortex. 26:4563–4573.

Huang YZ, Edwards MJ, Rounis E, Bhatia KP, Rothwell JC. 2005. Theta burst stimulation of the human motor cortex. Neuron. 45:201–206.

Hwang JH, Kim SH, Park CS, Bang SA, Kim SE. 2010. Acute high-frequency rTMS of the left dorsolateral prefrontal cortex and attentional control in healthy young men. Brain Res. 1329:152–158.

Jackson RL, Ralph MAL, Pobric G. 2015. The Timing of Anterior Temporal Lobe Involvement in Semantic Processing. Journal of Cognitive Neuroscience. 27:1388–1396.

Jung J, Lambon Ralph MA. 2016. Mapping the Dynamic Network Interactions Underpinning Cognition: A cTBS-fMRI Study of the Flexible Adaptive Neural System for Semantics. Cereb Cortex. 26:3580–3590.

Kay G. 1991. Repeated Testing Applications Employing Computer-Based Performance Assessment Measures. J Clin Exp Neuropsyc. 13:50–50.

Lambon Ralph MA. 2014. Neurocognitive insights on conceptual knowledge and its breakdown. Philos Trans R Soc Lond B Biol Sci. 369:20120392.

Lambon Ralph MA, Jefferies E, Patterson K, Rogers TT. 2017. The neural and computational bases of semantic cognition. Nat Rev Neurosci. 18:42–55.

Lambon Ralph MA, Pobric G, Jefferies E. 2009. Conceptual knowledge is underpinned by the temporal pole bilaterally: convergent evidence from rTMS. Cereb Cortex. 19:832–838.

Lambon Ralph MA, Sage K, Jones RW, Mayberry EJ. 2010. Coherent concepts are computed in the anterior temporal lobes. P Natl Acad Sci USA. 107:2717–2722.

Lenz M, Galanis C, Muller-Dahlhaus F, Opitz A, Wierenga CJ, Szabo G, Ziemann U, Deller T, Funke K, Vlachos A. 2016. Repetitive magnetic stimulation induces plasticity of inhibitory synapses. Nat Commun. 7:10020.

Lisanby SH, Gutman D, Luber B, Schroeder C, Sackeim HA. 2001. Sham TMS: intracerebral measurement of the induced electrical field and the induction of motor-evoked potentials. Biol Psychiatry. 49:460–463.

Luber B, Lisanby SH. 2014. Enhancement of human cognitive performance using transcranial magnetic stimulation (TMS). Neuroimage. 85 Pt 3:961–970.

Miniussi C, Rossini PM. 2011. Transcranial magnetic stimulation in cognitive rehabilitation. Neuropsychol Rehabil. 21:579–601.

Mollo G, Cornelissen PL, Millman RE, Ellis AW, Jefferies E. 2017. Oscillatory Dynamics Supporting Semantic Cognition: MEG Evidence for the Contribution of the Anterior Temporal Lobe Hub and Modality-Specific Spokes. PLoS One. 12:e0169269.

Muller-Dahlhaus F, Vlachos A. 2013. Unraveling the cellular and molecular mechanisms of repetitive magnetic stimulation. Front Mol Neurosci. 6:50.

Mummery CJ, Patterson K, Price CJ, Ashburner J, Frackowiak RS, Hodges JR. 2000. A voxel-based morphometry study of semantic dementia: relationship between temporal lobe atrophy and semantic memory. Ann Neurol. 47:36–45.

Murphy C, Rueschemeyer SA, Watson D, Karapanagiotidis T, Smallwood J, Jefferies E. 2017. Fractionating the anterior temporal lobe: MVPA reveals differential responses to input and conceptual modality. Neuroimage. 147:19–31.

O’Shea J, Johansen-Berg H, Trief D, Gobel S, Rushworth MF. 2007. Functionally specific reorganization in human premotor cortex. Neuron. 54:479–490.

Oldfield RC. 1971. The assessment and analysis of handedness: the Edinburgh inventory. Neuropsychologia. 9:97–113.

Pascual-Leone A, Valls-Sole J, Wassermann EM, Hallett M. 1994. Responses to rapid-rate transcranial magnetic stimulation of the human motor cortex. Brain. 117 (Pt 4):847–858.

Patterson K, Nestor PJ, Rogers TT. 2007. Where do you know what you know? The representation of semantic knowledge in the human brain. Nat Rev Neurosci. 8:976–987.

Peelen MV, Caramazza A. 2012. Conceptual object representations in human anterior temporal cortex. J Neurosci. 32:15728–15736.

Pell GS, Roth Y, Zangen A. 2011. Modulation of cortical excitability induced by repetitive transcranial magnetic stimulation: influence of timing and geometrical parameters and underlying mechanisms. Prog Neurobiol. 93:59–98.

Pobric G, Jefferies E, Lambon Ralph MA. 2010. Category-specific versus category-general semantic impairment induced by transcranial magnetic stimulation. Curr Biol. 20:964–968.

Pobric G, Jefferies E, Ralph MA. 2007. Anterior temporal lobes mediate semantic representation: mimicking semantic dementia by using rTMS in normal participants. Proc Natl Acad Sci USA. 104:20137–20141.

Pobric G, Jefferies E, Ralph MA. 2010. Amodal semantic representations depend on both anterior temporal lobes: evidence from repetitive transcranial magnetic stimulation. Neuropsychologia. 48:1336–1342.

Pobric G, Lambon Ralph MA, Jefferies E. 2009. The role of the anterior temporal lobes in the comprehension of concrete and abstract words: rTMS evidence. Cortex. 45:1104–1110.

Ragert P, Dinse HR, Pleger B, Wilimzig C, Frombach E, Schwenkreis P, Tegenthoff M. 2003. Combination of 5 Hz repetitive transcranial magnetic stimulation (rTMS) and tactile coactivation boosts tactile discrimination in humans. Neurosci Lett. 348:105–108.

Ralph MA, Jefferies E, Patterson K, Rogers TT. 2017. The neural and computational bases of semantic cognition. Nat Rev Neurosci. 18:42–55.

Rice GE, Hoffman P, Lambon Ralph MA. 2015. Graded specialization within and between the anterior temporal lobes. Ann N Y Acad Sci.

Ridding MC, Ziemann U. 2010. Determinants of the induction of cortical plasticity by non-invasive brain stimulation in healthy subjects. J Physiol-London. 588:2291–2304.

Rogers TT, Patterson K, Jefferies E, Ralph MA. 2015. Disorders of representation and control in semantic cognition: Effects of familiarity, typicality, and specificity. Neuropsychologia. 76:220–239.

Rossi S, Rossini PM. 2004. TMS in cognitive plasticity and the potential for rehabilitation. Trends Cogn Sci. 8:273–279.

Shimotake A, Matsumoto R, Ueno T, Kunieda T, Saito S, Hoffman P, Kikuchi T, Fukuyama H, Miyamoto S, Takahashi R, Ikeda A, Lambon Ralph MA. 2015. Direct Exploration of the Role of the Ventral Anterior Temporal Lobe in Semantic Memory: Cortical Stimulation and Local Field Potential Evidence From Subdural Grid Electrodes. Cereb Cortex. 25:3802–3817.

Sole-Padulles C, Bartres-Faz D, Junque C, Clemente IC, Molinuevo JL, Bargallo N, Sanchez-Aldeguer J, Bosch B, Falcon C, Valls-Sole J. 2006. Repetitive transcranial magnetic stimulation effects on brain function and cognition among elders with memory dysfunction. A randomized sham-controlled study. Cereb Cortex. 16:1487–1493.

Terao Y, Ugawa Y, Suzuki M, Sakai K, Hanajima R, Gemba-Shimizu K, Kanazawa I. 1997. Shortening of simple reaction time by peripheral electrical and submotor-threshold magnetic cortical stimulation. Exp Brain Res. 115:541–545.

Vallar G, Bolognini N. 2011. Behavioural facilitation following brain stimulation: implications for neurorehabilitation. Neuropsychol Rehabil. 21:618–649.

Visser M, Jefferies E, Embleton KV, Lambon Ralph MA. 2012. Both the middle temporal gyrus and the ventral anterior temporal area are crucial for multimodal semantic processing: distortion-corrected fMRI evidence for a double gradient of information convergence in the temporal lobes. J Cogn Neurosci. 24:1766–1778.

Visser M, Jefferies E, Lambon Ralph MA. 2010. Semantic processing in the anterior temporal lobes: a meta-analysis of the functional neuroimaging literature. J Cogn Neurosci. 22:1083–1094.

Vlachos A, Muller-Dahlhaus F, Rosskopp J, Lenz M, Ziemann U, Deller T. 2012. Repetitive magnetic stimulation induces functional and structural plasticity of excitatory postsynapses in mouse organotypic hippocampal slice cultures. J Neurosci. 32:17514–17523.

Wagner M, Rihs TA, Mosimann UP, Fisch HU, Schlaepfer TE. 2006. Repetitive transcranial magnetic stimulation of the dorsolateral prefrontal cortex affects divided attention immediately after cessation of stimulation. J Psychiatr Res. 40:315–321.

Wassermann EM, Epstein CM, Ziemann U. 2008. The Oxford handbook of transcranial stimulation. Oxford; New York: Oxford University Press.

