## Supplemental information for "Enhancing vs. inhibiting semantic performance with repetitive transcranial magnetic stimulation over the anterior temporal lobe: frequency- and task- specific effects"

**Supplementary Information**

Supplementary Table 1

Supplementary Figure 1

Supplementary Figure 2

Supplementary Figure 3

**Supplementary Table 1**

| Items | Category | Items | Category | Items | Category |
| --- | --- | --- | --- | --- | --- |
| collie | dog | gorilla | primates | spoon | Kitchen utensils |
| alsatian | dog | chimpanzee | primates | whisk | Kitchen utensils |
| dalmatian | dog | baboon | primates | peeler | Kitchen utensils |
| poodle | dog | marmoset | primates | spatula | Kitchen utensils |
| pekinese | dog | orang-utan | primates | peppermill | Kitchen utensils |
| bulldog | dog | rat | rodents | garlic press | Kitchen utensils |
| greyhound | dog | mouse | rodents | pestle & mortar | Kitchen utensils |
| robin | birds | squirrel | rodents | palette knife | Kitchen utensils |
| swan | birds | hamster | rodents | runner bean | legume |
| pigeon | birds | mole | rodents | pea | legume |
| owl | birds | porcupine | rodents | mange toute | legume |
| kingfisher | birds | raccoon | rodents | french bean | legume |
| pheasant | birds | Ford | cars | courgette | legume |
| heron | birds | Jaguar | cars | aubergine | legume |
| wren | birds | Rolls Royce | cars | apple | fruit |
| lion | cats | Ferrari | cars | pear | fruit |
| tiger | cats | BMW | cars | blackberry | fruit |
| leopard | cats | Boeing 747 | aeroplanes | blueberry | fruit |
| cheetah | cats | Spitfire | aeroplanes | raspberry | fruit |
| jaguar | cats | Lear | aeroplanes | strawberry | fruit |
| panther | cats | Concorde | aeroplanes | gooseberry | fruit |
| lynx | cats | Hercules | aeroplanes | apricot | fruit |
| python | snakes | AWAX | aeroplanes | oak | trees |
| adder | snakes | trilby | hats | horse chesnut | trees |
| anaconda | snakes | deerstalker | hats | maple | trees |
| shark | fish | panama | hats | fir | trees |
| trout | fish | stetson | hats | willow | trees |
| salmon | fish | fez | hats | cherry | trees |
| plaice | fish | mitre | hats | daisy | flowers |
| ray | fish | wellington's | footwear | daffodil | flowers |
| tuna | fish | trainers | footwear | tulip | flowers |
| Friesian | cows | slippers | footwear | rose | flowers |
| Jersey | cows | stilettos | footwear | bluebell | flowers |
| Aberdeen Angus | cows | doc martin's | footwear | lily | flowers |
| lobster | shellfish | brogues | footwear | baguette | bread |
| crab | shellfish | spade | Garden tools | croissant | bread |
| octopus | shellfish | fork | Garden tools | bloomer | bread |
| jellyfish | shellfish | secateurs | Garden tools | crumpet | bread |
| squid | shellfish | shears | Garden tools | pikelet | bread |
| oyster | shellfish | hoe | Garden tools | cheddar | cheese |
| clam | shellfish | trowel | Garden tools | double gloucester | cheese |
|  |  | parmesan | cheese | brie | cheese |
|  |  | edam | cheese | stilton | cheese |

Table S1. The list of items for category judgement task

**Supplementary Figure 1**

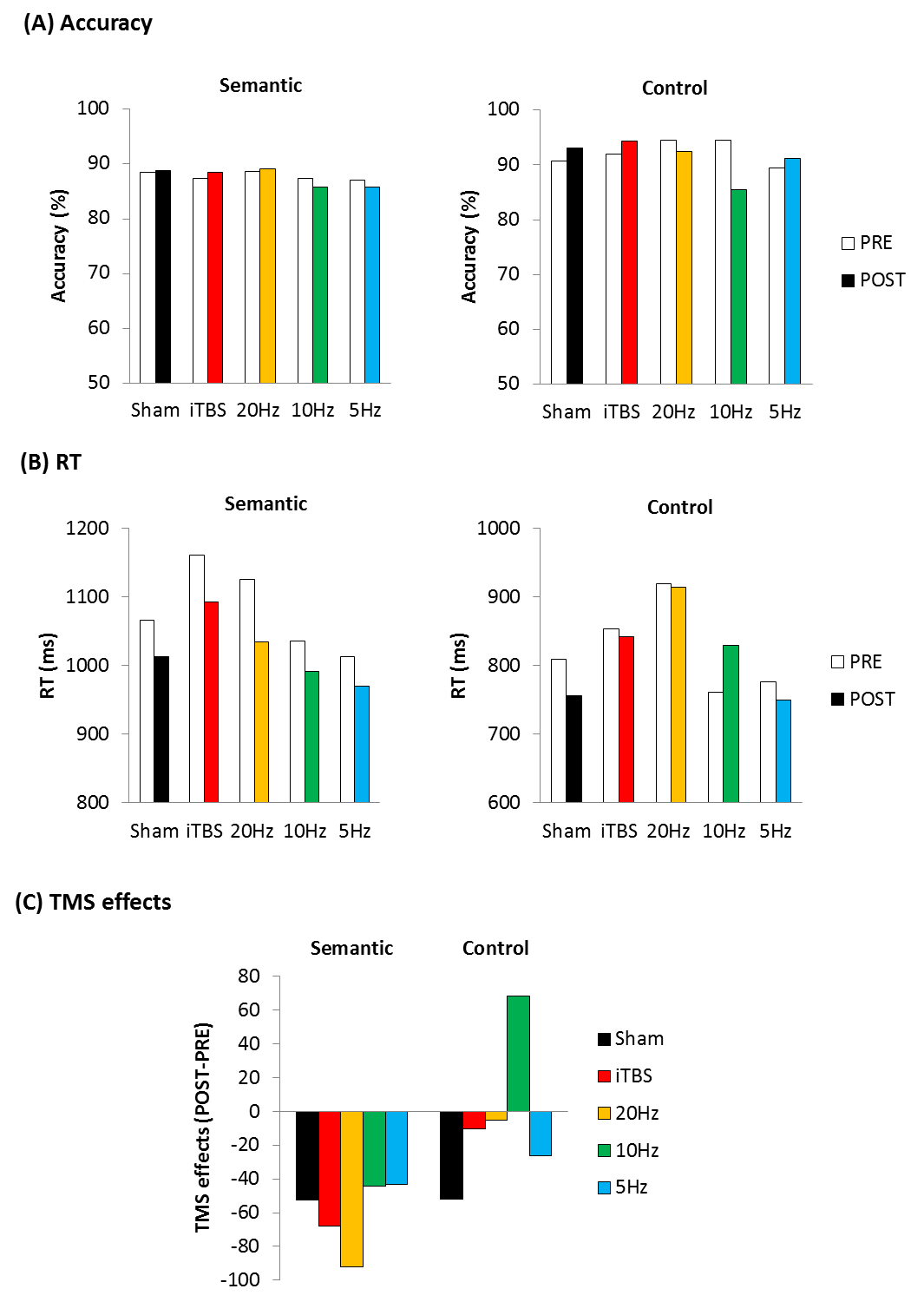

Fig S1 The results of the pilot study. (A) The averaged accuracy for the category judgment task (left) and control task (right). (B) The averaged reaction time (RT) for the category judgment task (left) and control task (right). (C) TMS effects were calculated by subtracting RT in PRE from POST TMS session. Negative value indicates the facilitaotry effects. 20Hz stimulation showed the facilitatory effects in the category judgment task compared to sham.

**Supplementary Figure 2**

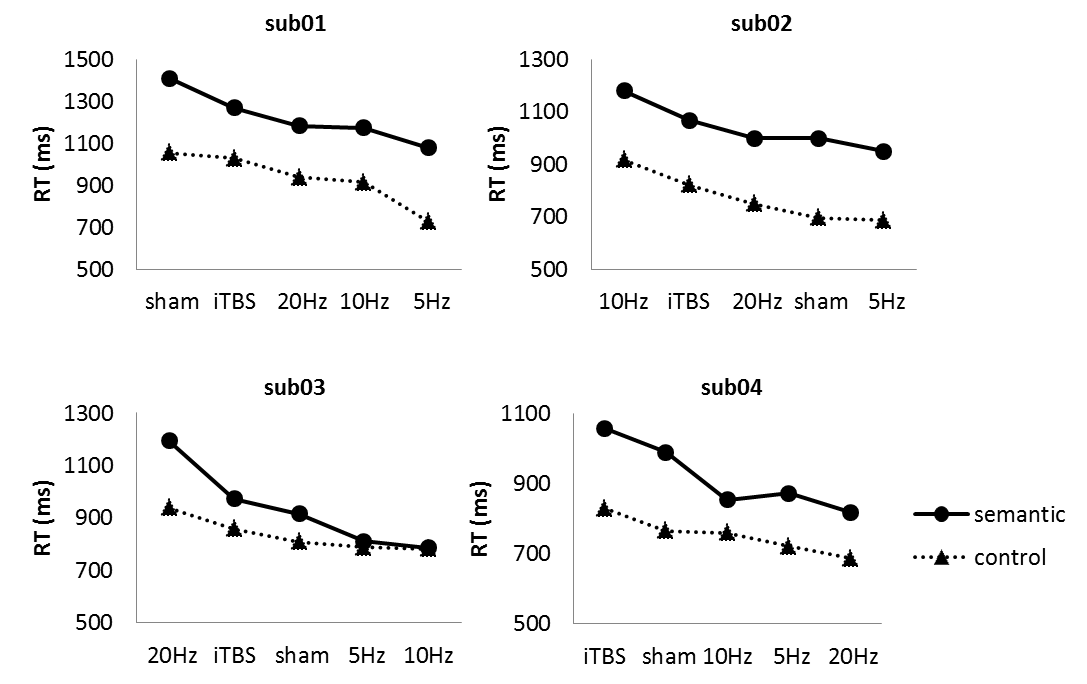

Fig S2. Individual task performance (RT) in the PRE TMS sessions. X-axis indicates the order of each protocol according to individuals (left-first session, right-last session). Regardless of the tasks, there were huge practice (order) effects.

**Supplementary Figure 3**

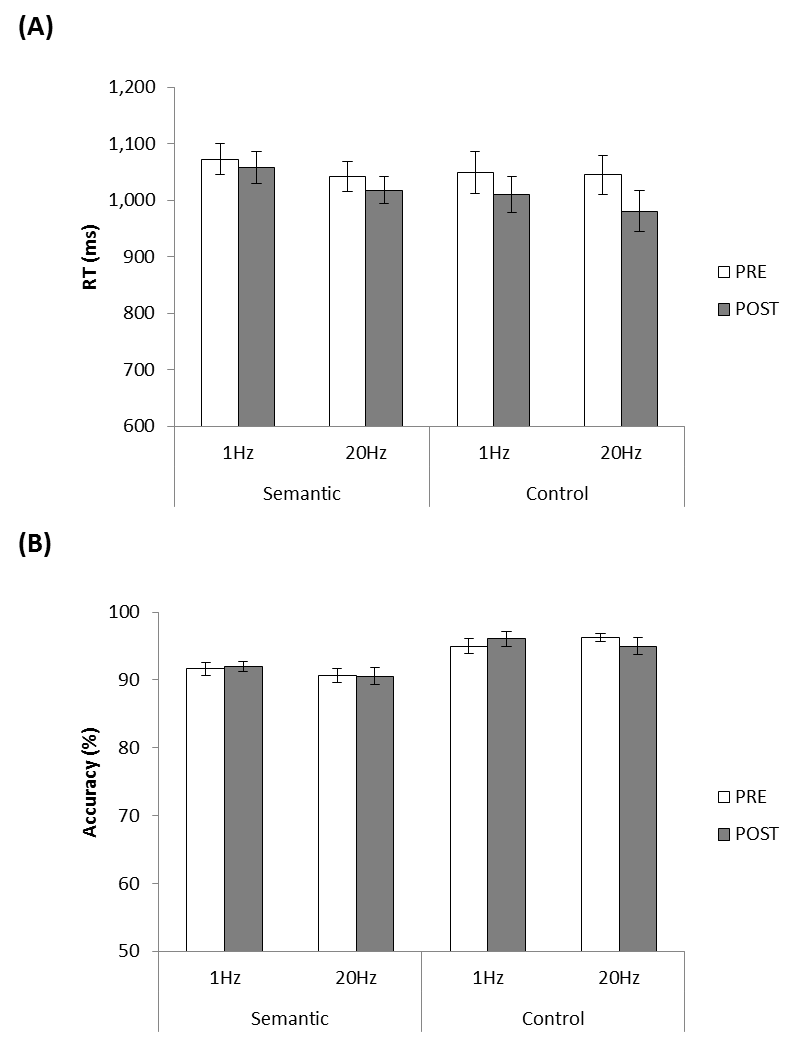

Fig S3. The results of sham stimulation. (A) The averaged RT for the category judgment task and control task. (B) The averaged accuracy for the category judgment task and control task.
